# Spatial and Compositional Biomarkers in Tumor Microenvironment Predicts Clinical Outcomes in Triple-Negative Breast Cancer

**DOI:** 10.1101/2023.12.18.572234

**Authors:** Haoyang Mi, Ravi Varadhan, Ashley M. Cimino-Mathews, Leisha A. Emens, Cesar A. Santa-Maria, Aleksander S. Popel

## Abstract

Triple-negative breast cancer (TNBC) is an aggressive subtype of breast cancer with limited treatment options, which warrants identification of novel therapeutic targets. Deciphering nuances in the tumor microenvironment (TME) may unveil insightful links between anti-tumor immunity and clinical outcomes, yet such connections remain underexplored. Here we employed a dataset derived from imaging mass cytometry of 58 TNBC patient specimens at single-cell resolution and performed in-depth quantifications with a suite of multi-scale computational algorithms. We detected distinct cell distribution patterns among clinical subgroups, potentially stemming from different infiltration related to tumor vasculature and fibroblast heterogeneity. Spatial analysis also identified ten recurrent cellular neighborhoods (CNs) - a collection of local TME characteristics with unique cell components. Coupling of the prevalence of pan-immune and perivasculature immune hotspot CNs, enrichment of inter-CN interactions was associated with improved survival. Using a deep learning model trained on engineered spatial data, we can with high accuracy (mean AUC of 5-fold cross-validation = 0.71) how a separate cohort of patients in the NeoTRIP clinical trial will respond to treatment based on baseline TME features. These data reinforce that the TME architecture is structured in cellular compositions, spatial organizations, vasculature biology, and molecular profiles, and suggest novel imaging-based biomarkers for treatment development in the context of TNBC.

## INTRODUCTION

Breast cancer is the most prevalent non-dermatologic cancer in women in the United States with 297,790 new cases and 43,170 deaths predicted in 2023^1^. Among all breast cancer subtypes, triple-negative breast cancer (TNBC) has the highest high risk of recurrence and mortality if metastatic. TNBC is characterized by the absence of expression of estrogen receptor (ER), progesterone receptor (PR) and human epidermal growth factor receptor-2 (HER2), making drugs that target these pathways in other subtypes not a viable option here. Despite limited benefits, chemotherapy remains the mainstay of systemic treatment for early- and late-stage TNBC^2^. Advances in immunotherapies that target immune checkpoint molecules have improved clinical outcomes for patients with solid tumors, including those with TNBC. Recently, a new standard-of-care was established when the US Food and Drug Administration (FDA) approved pembrolizumab, a programmed cell death protein 1 (PD-1) inhibitor, combined with chemotherapy for high-risk early stage TNBC as neoadjuvant treatment, and then continued as adjuvant monotherapy following tumor resection^3^. Pembrolizumab has also been approved in the metastatic setting for patients who have PD-L1 positive (CPS ≥ 10) TNBC; many types of immunotherapy and combinations are also under investigtation^4–7^. Although both in the early-stage and metastatic setting the addition of pembrolizumab has improved outcomes in some patients, the majority of patients do not benefit and responders may eventually recur. Thus, a complete landscape of tumor progression and antitumor immunity remains unclear and warrants urgent investigations.

The complexity and heterogeneity of the tumor microenvironment (TME) is a hallmark of resistance to therapy^8^. The distributions and spatial interplay of cell populations is known to inform the overall efficacy of tumor control^9–11^; therefore, dissecting the TME at single-cell resolution may reveal clinically relevant tissue features, provide mechanistic insights into antitumor immunity, and identify biomarkers for novel treatment design. Development of multiplexed imaging technologies have now enabled deep *in situ* profiling of TME components simultaneously, thereby facilitating our understanding of the nuances in TNBC. For instance, Keren et al have revealed the prognostic value of an ordered immune structure along with expression of checkpoint molecules using multiplexed ion beam imaging^12^. Recently, Danenberg et al used imaging mass cytometry (IMC) to generate 37-dimensional images of tumors from 718 breast cancer patients in the METABRIC study, and this dataset represents a valuable resource for the breast cancer research community^13^. Despite these advances, the links between clinical outcomes and TME contextures in TNBC are still incompletely understood. For example, recent literature presents conflicting evidence on the prognostic value of B cells. While B cells have been correlated with FoxP3^+^ regulatory T cells and associated with poor survival, they also have been linked to favorable prognosis via IgG-mediated humoral immunity^14,15^. In addition, computational approaches, especially spatial analysis methods, have lagged behind technological innovations. To date, these approaches primarily focus on quantifying simplistic (albeit important) features such as nearest neighbor distance, which fail to recapitulate the complexity of the TME. Adding to the challenge, studies are often restricted to a single cohort due to cost and patient recruitment difficulties, thereby constraining extrapolation to a broader context.

To complement ongoing endeavors in mapping the complete landscape of the TME and connections to clinical outcomes in TNBC, we acquired open-source TNBC datasets at single-cell resolution^13,16^ derived from multiplexed imaging systems, and leveraged a computational pipeline with a suite of image analysis algorithms. With a discovery cohort of 58 TNBC patients from the METABRIC dataset, we report substantial intra-tumoral heterogeneity of cell populations, tumor vasculature, and tissue architecture that were associated with clinical features. Candidate biomarkers were distilled from such differential spatial and compositional TME features. By integrating these data with deep learning, we were able to predict with high accuracy the treatment response of a separate cohort of TNBC patients in the NeoTRIP clinical trial^17^. The NeoTRIP study randomized patients with TNBC to receive neoadjuvant carboplatin and nab-paclitaxel (chemotherapy) with or without atezolizumab (a PD-L1 inhibitor). Collectively, our study provides a framework to apply multiplexed imaging analysis for biomarker discovery and adds an important piece towards comprehending the single-cell landscape of TNBC.

## RESULTS

### Patient selection and data acquisition

To develop and leverage quantitative methods for an in-depth interrogation of the TME in TNBC, we utilized a dataset with single-cell resolution comprised of 718 breast cancer patients previously enrolled in the METABRIC study^13,18,19^. Among these patients, we selected a cohort of 58 patients with TNBC for whom clinical information was also available for downstream analysis (**Fig. 1A**). Briefly, tissue microarray (TMA) slides containing breast tumor samples were labeled with a panel of 35 antibodies and analyzed using IMC to generate multiplexed images. In cases where multiple regions were taken from the same patient (*n* = 6), one was randomly selected to represent that individual. Majority of selected patients (*n* = 51) have profiled regions (tissue cores) with a diameter of ~500 µm. The remaining patients (*n* = 7) have profiled regions with a diameter of 1 mm. An image processing pipeline was then followed to discriminate tissue compartments (tumor region and vasculature) and segment and phenotype single cells (**Fig. 1B** and ref.^13^). This workflow results in a collection of 131,001 cells of 15 phenotypes that include lymphoid, myeloid, stromal, and epithelial (tumor) lineages (**Fig. 1C, D**). For biomarker validation and deep learning, we utilized a single-cell dataset collected from the NeoTRIP clinical trial. The two cohorts share the majority of technical details such as imaging acquisition, segmentations and phenotyping, and therefore represent an ideal resource for comparative analysis. The discovery cohort (METABRIC cohort) was subjected to evaluation of associations between TME features and downstream clinical outcomes.

**Figure 1.**
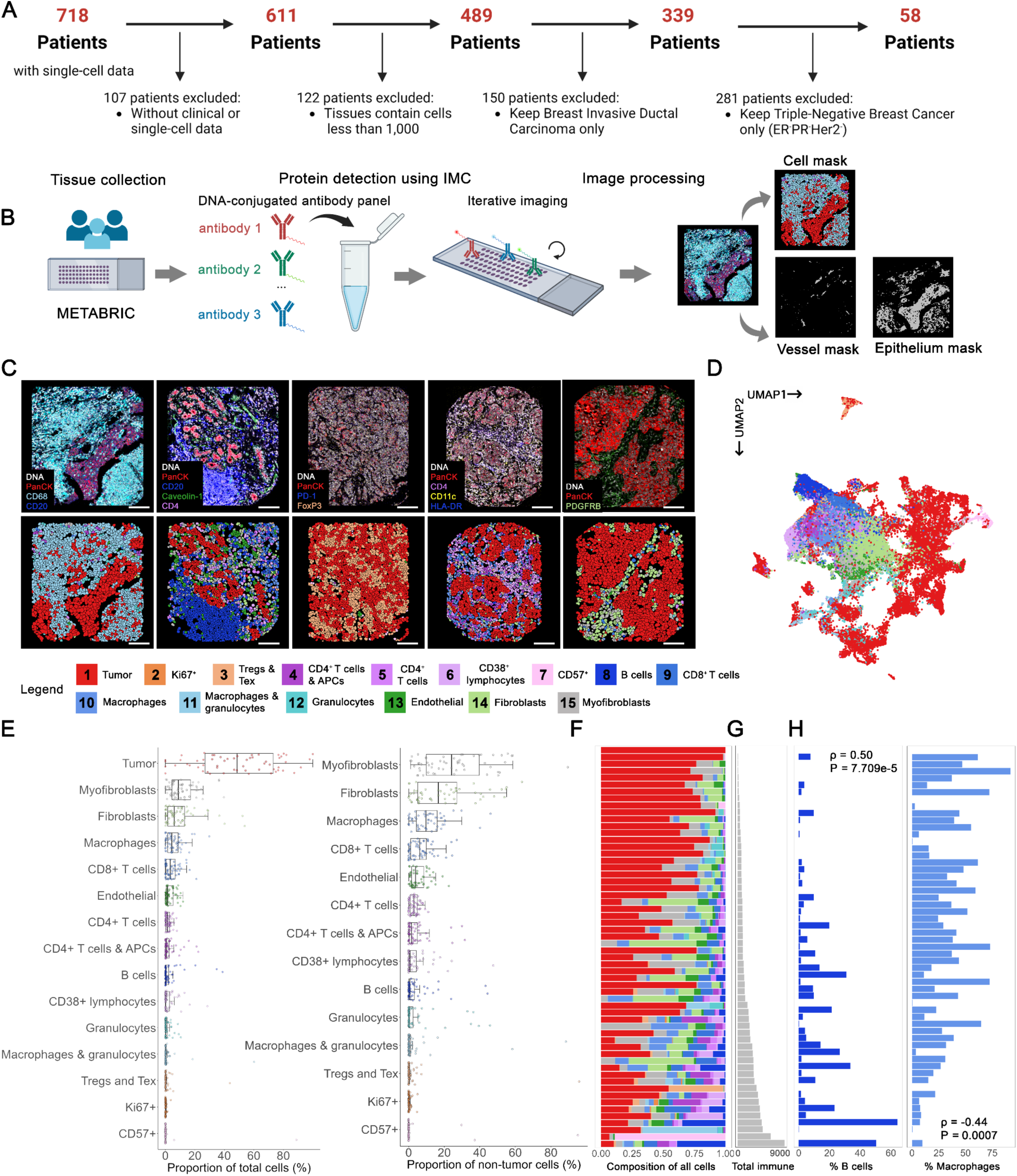
Multiplexed tissue imaging enables in situ single-cell profiling of the TME architectures. **(A)** Patient selection criteria: patients with both clinical and single-cell data available; cores in tissue microarray contained at least 1,000 cells for effective characterization of tumor-immune landscape; breast invasive ductal adenocarcinoma; triple-negative subtype. **(B)** Description of materials. The discovery cohort was collected from METABRIC study. Samples were imaged using image mass cytometry and processed to generate a single-cell dataset and tissue masks. **(C)** Representative TMA cores depicted as three-color images to illustrate tissue dynamics with 15 identified cell types. **(D)** Uniform Manifold Approximation and Projection (UMAP) revealed TME heterogeneity in cellular compositions. **(E)** Prevalence of 15 cell types, including 14 non-tumor cell types, across 58 patients with TNBC as a proportion of total cells and non-tumor cells. **(F)-(H)**: **(F)** Waterfall stacked bar plots describing the distribution of 15 cell types per patient and **(G)** ranked in the ascending order of immune cell density. **(H)** Proportion of B cells increases whereas proportion of macrophages decreases with immune cell density.

### Clinical subgrouping

Patients were categorized based upon the following clinicopathologic features: age at diagnosis, relative to study median age (≤ 55 versus > 55 years of age); tumor cellularity (low versus moderate versus high); tumor histologic grade (1-2 versus 3); tumor stage (I-II versus III-IV); primary tumor laterality (left versus right); relapse status at the time of last follow-up (recurred versus not recurred); tumor mutational burden (TMB), relative to study median (≤ 5, denoted as low versus > 5, denoted high); overall survival (OS), relative to study median (≤ 66 versus > 66 months), with a mean OS of 33 months in short-term survivors and a mean OS of 157 months in long-term survivors. Survival differences for each clinical subgroups were assessed using Kaplan-Meier plots (**Supplementary Figure S1)**.

### Clinical significance of single-cell distributions in the TNBC tumor microenvironment

We reasoned that the TME structure is a key window into antitumor immunity. Accordingly, we quantified the prevalence of each cell population across patients and observed remarkable variability (**Fig. 1E**). Tumor cells were the most prevalent type (mean proportion = 48.6%), and myofibroblasts (mean proportion = 25.8%) were the most abundant non-tumor cell type, followed by macrophages (mean proportion = 11.1%) and CD8^+^ T cells (mean proportion = 7.1%) as the primary myeloid and lymphoid cell populations, respectively (**Fig. 1F**). The TME compositions also varied across patients, with distinct immune infiltration profiles spanning from tumor-dominant to immune-dominant (**Fig. 1F, G**). Interestingly, patients with more immune cells tended to have a higher fraction of B cells (Pearson R = 0.50, *p* < 10^-4^) and a lower fraction of macrophages (Pearson R = 0.44, *p* = 0.0007) (**Fig. 1H, Supplementary Figure S2A**). Assuming the level of immune infiltration models the TME over time, these results may indicate a synchronization of immune cell subsets entering the tumor. To further gauge the ordering in immune infiltration, we analyzed the co-occurrence of immune populations across patients. We classified different immune populations for each patient as present or absent if there was at least one cell count. Then we applied chi-square test to evaluate the co-occurrence pattern (**Supplementary Figure S2B**). We found notable correlations between immune populations across patients. For example, patients that had macrophages also had B cells, CD8^+^ T cells and CD4^+^ T cells (χ^2^ *p* < 0.01, *p* < 0.0001, *p* < 0.00001 respectively), which were in line with our previous study that evaluated the correlations between CD4, CD8 and CD20 markers^20^. Altogether, these results support a notion that tumor microenvironment was coordinated by specific immune cells in a location-specific, context-dependent manner.

To further assess the associations between cellular distribution patterns and clinical outcomes, we quantified the number of cell types for each patient, and compared clinically relevant subgroups of patients. Indeed, we found significant heterogeneity within and across clinical subgroups (**Supplementary Fig. S3**). These differences may further be mitigated or intensified due to the antagonistic interaction between cells. For example, high ratios of CD4^+^ T cells to macrophages were significantly associated with lower pathologic stages; high ratios of a T cell subset (collection of cell types 4, 5, 9) to fibroblasts were significantly associated with patients who had not relapsed (**Fig. 2A, B**); and none of these cell types was independently linked to the respective outcomes (**Fig. 2C**). As expected, elevated fractions of CD4^+^ and CD8^+^ T cells, indicative of an immune-activated environment, were associated with improved survival^21–23^. Although past reports have widely described the immunosuppressive role of fibroblasts in breast cancer, we found enrichment of fibroblasts in long-term survivors (**Fig. 2D**). To reconcile this contradiction, we set out to further dissect fibroblasts into four subpopulations with unsupervised hierarchical clustering (**Fig. 2D, Supplementary Figures 4A-B**). Subclustering showed that fibroblasts were characterized by heterogeneous protein expression, which were defined as following: Fibro-C1 (SMA^Low-Med^ Caveolin-1^Hi^ PDGFRB^Hi^ FSP1^Low-Med^); Fibro-C2 (SMA^Med-Hi^ Caveolin-1^Med^ PDGFRB^Med^ FSP1^Med^); Fibro-C3 (SMA^Hi^ Caveolin-1^Low^ PDGFRB^Low^ FSP1^Neg-Low^); and Fibro-C4 (SMA^Neg-Low^ Caveolin-1^Neg-Low^ PDGFRB^Low^ FSP1^Hi^) (**Fig. 2E, Supplementary Figure 4C**). We found notable differences between the compositions of these clusters among survival subgroups, marked by the enrichment of Fibro-C3 in long-term survivors (**Fig. 2F-G**). This complements a recent finding that a subset of fibroblasts (noted as CAF-S4) with marker expression profiles closely resembling Fibro-C3 was associated with decreased risks of inhibition on effector T cells (albeit not linked to survival benefits)^24,25^. Of note, we reported that Fibro-C3 had a unique negative correlation trend with B cells and tumor cell counts, which may indicate a novel mechanism of anti-tumor immunity mediated by a specific B cell-fibroblast interacting axis (**Supplementary Figure S4D**).

**Figure 2.**
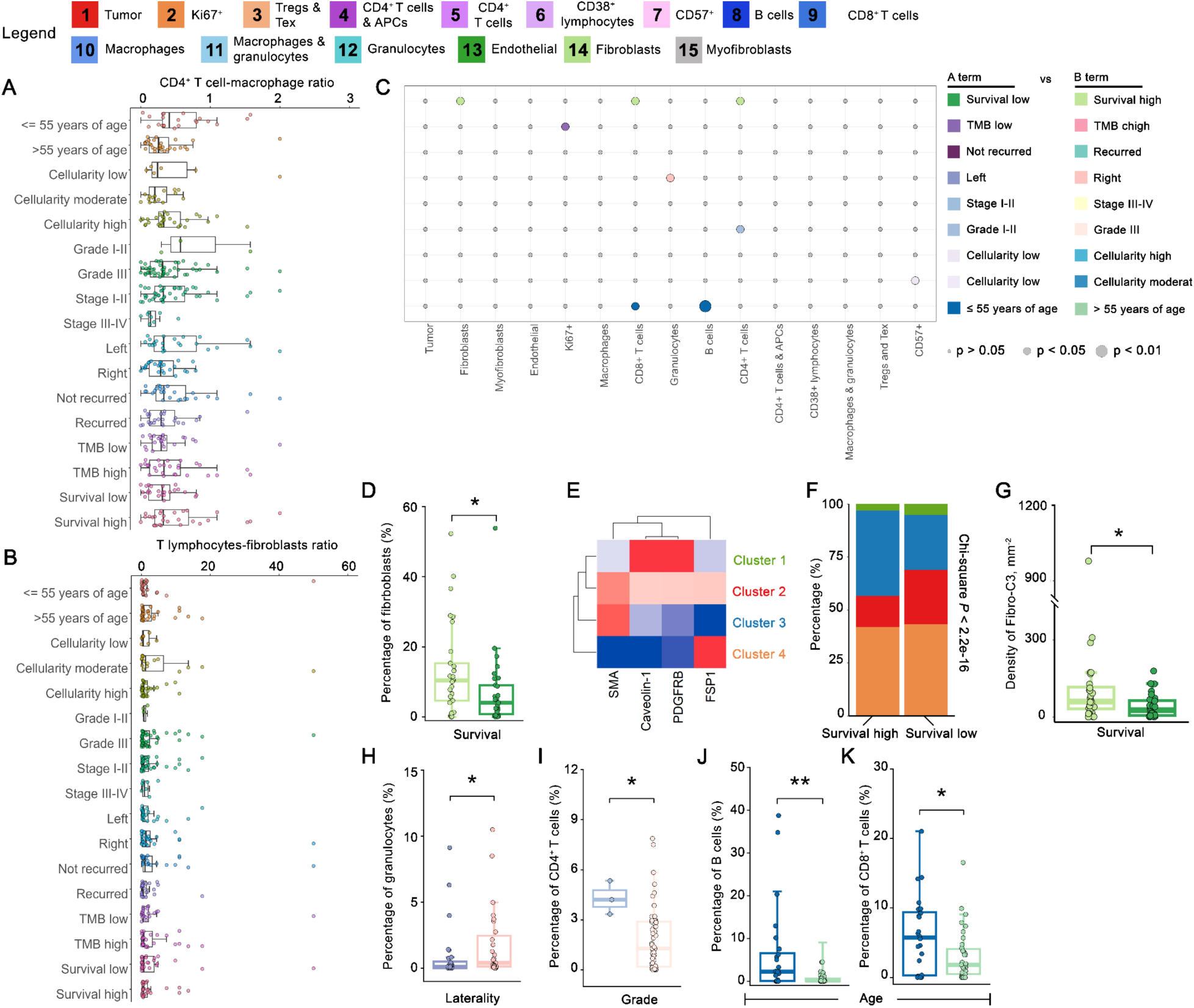
Heterogeneous single-cell distributions across clinico-pathological features in TNBC. **(A)** Distributions of CD4^+^ T cell – macrophage ratios across age at diagnosis (55 years of age or younger *n* = 24, older than 55 years of age *n* = 34), cellularity (low *n* = 8, moderate *n* = 16, high *n* = 32), grade (I-II *n* = 3, III *n* = 55), stage (I-II *n* = 47, III-IV *n* = 8), primary tumor laterality (left *n* = 28, right *n* = 27), relapse-free status (recurred *n* = 28, not recurred *n* = 30), TMB (low *n* = 25, high *n* = 33), overall survival (short-term *n* = 29, long-term *n* = 29, median thresholding). **(B)** Distribution of T lymphocyte – fibroblast ratios across clinical subgroups. **(C)** Clinical significance of cell distributions across clinical subgroups depicted using bubble plot, in which the circle color represents which of the two comparisons (A term or B term) on the *y* axis has higher proportions of the cell type on the *x* axis and circle size represents the level of significance. **(D)** Prevalence of fibroblasts in long-term and short-term survivors as a proportion of total cells. **(E)** Hierarchical clustering on four fibroblast markers (SMA, Caveolin-1, PDGFRβ and FSP1) revealed four subclusters with unique marker expression profiles. **(F)** Compositions of fibroblast subclusters in long-term and short-term survivors. **(G)** Prevalence of Fibro-C3 in long-term and short-term survivors as densities. **(H)** Prevalence of granulocytes by primary tumor laterality as percentages of total cells. **(I)** Prevalence of CD4^+^ T cells by TNBC grade categories as percentages of total cells. Prevalence of **(J)** B cells and **(K)** CD8^+^ T cells by age categories as percentages of total cells. Statistical comparisons, either paired or unpaired, were performed using Wilcoxon rank-sum test. **, *p* < 0.01; *, *p* < 0.05.

In addition, we reported higher frequencies of granulocytes in the right-sided TNBC (**Fig. 2H**; Wilcoxon rank-sum *p* = 0.047) and higher frequencies of CD4^+^ T cells in lower TNBC grade (**Fig. 2I**; Wilcoxon rank-sum *p* = 0.02), although such slight difference maybe likely due to small sample size of the discovery cohort. Interestingly, B cells and CD8^+^ T cells accumulated in an age-dependent manner (**Fig. 2J, K**; Wilcoxon rank-sum *p* = 0.0084 and 0.045, respectively), which may infer a significant decline in the infiltration of effective T cells in the TME of older patients.

### Vascular density predict relapse-free survival (RFS)

Given the importance of fibroblasts in tumor angiogenesis^26,27^, we further assessed the relationship between TNBC vascular architecture and clinical outcomes. Here, vasculatures were assessed across patients by a vessel classifier that was developed based on CD31, SMA and Caveolin-1 expressions (see ref.^13^ and **Methods and Materials**). Detected vessels larger than 500 µm^2^ were denoted as valid vessels and selected for downstream analysis to minimize the effect of staining artifacts and mis-classification. Cells within vessels were considered to have low impact on modulating local immune response since they had not extravasated and were therefore not included in the single-cell analysis. Distances between the remaining cells to their nearest valid vessel were computed to prepare for the spatial analysis (**Fig. 3A**).

**Figure 3.**
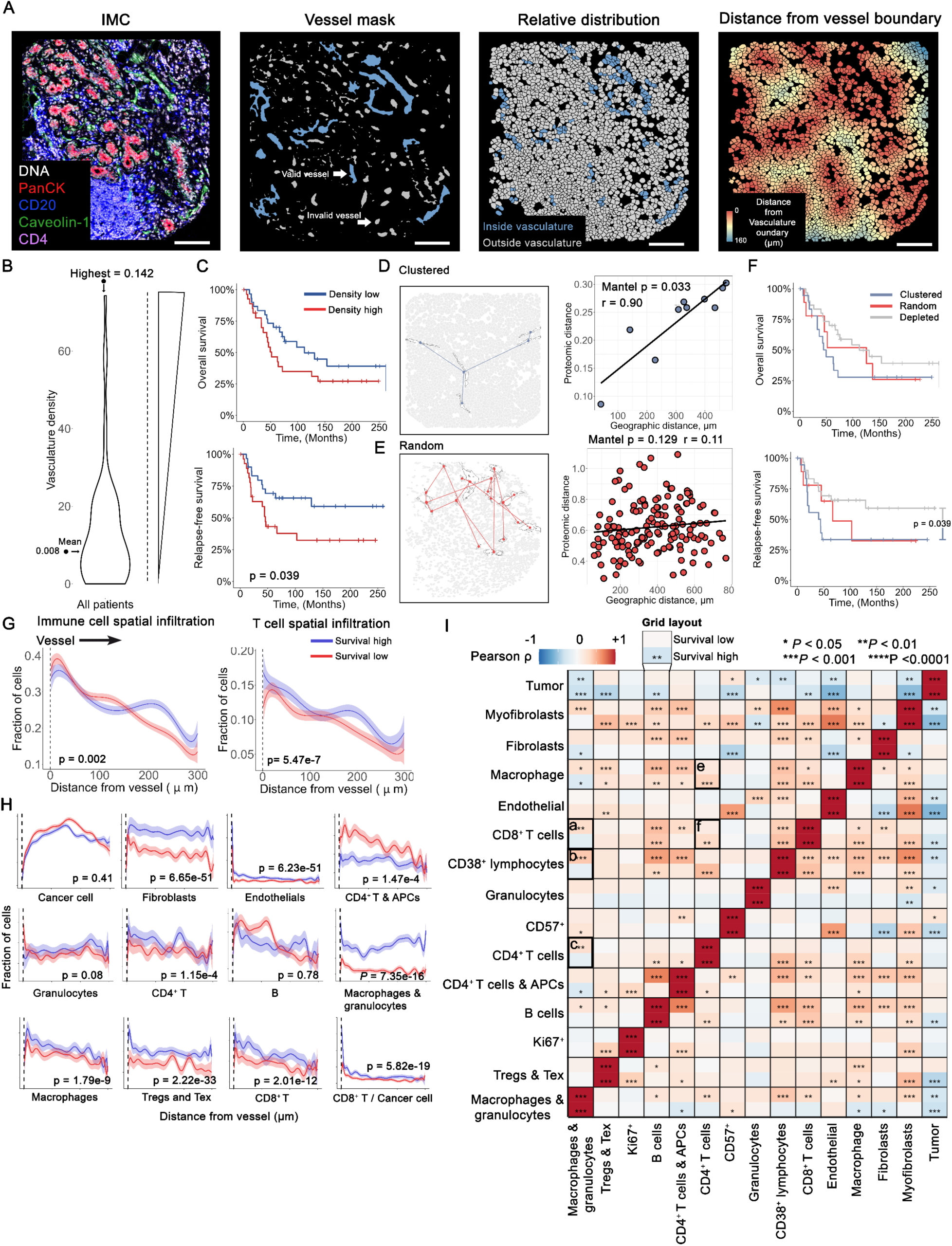
Multi-scale quantifications of perivascular niche provide insights into clinical significances. **(A)** Analytical workflow for generating valid vessel mask and valid cell set. **(B)** Prevalence of valid vessels (number of valid vessels per μm^2^) across all patients in discovery cohort. **(C)** Overall and RFS plots for valid vessel densities in TME where patients were classified into low or high density relative to the median across all patients. **(D)** Illustration of a TMA core with clustered perivascular niches (Patient MB-0350). Left panel: spatial locations of valid vessels and their nearest neighbors in molecular domain. Right panel: scatterplot of spatial and molecular distances. Each circle represents a pairwise distance between two valid vessels. R, Mantel correlation coefficient; p, Mantel test p value. **(E)** Illustration of a TMA core with random perivascular niches (Patient MB-0206). Left panel: spatial locations of valid vessels and their nearest neighbors in molecular domain. Right panel: scatterplot of spatial and molecular distances. Each circle represents a pairwise distance between two valid vessels. R, Mantel correlation coefficient; p, Mantel test p value. **(F)** Overall and RFS plots for patients with clustered perivascular niches, random perivascular niches, and valid vessel depleted patterns. Patients with no valid vessel had significantly superior RFS compared to patients with random pattern. Similar trend was observed in OS. **(G)** Infiltration profiles for immune cell and T lymphocyte proportions (fractions to total cells) as a function of distance to nearest vessel boundaries evaluated at various intervals. **(H)** Breakdown of infiltration profiles into each cell population. **(I)** Heatmap describing positive (red) or negative (blue) correlations between pairwise cell infiltration profiles across short-term (up grids) and long-term (bottom grids) survivors. Black boxes indicate associations referenced in the text. Wilcoxon rank-sum test was used for statistical tests and p-values were adjusted using Bonferroni method. ****, *p* < 0.0001; ***, *p* < 0.001; **, *p* < 0.01; *, *p* < 0.05.

To evaluate the prognostic significance of the vascular distributions, we computed vascular density (VD) across the discovery cohort and stratified patients into groups with low versus high vascular density using the mean (7.07 µm^-2^) as the threshold (**Fig. 3B** and **Supplementary Figure S5A**). While the distinct VD groups were not associated with significant differences in OS, the high VD group demonstrated notably worse RFS. This finding was consistent despite changing the area threshold across vessel masks from 300 to 700 µm^2^ (**Fig. 3C** and **Supplementary Figure S5B-C**). This finding reinforces the notion that increased VD may associated with higher risk of disease recurrence and metastasis. This correlation has been reported in various cancer types, including hepatocellular carcinoma and breast cancer^28–32^.

### STAMPER identifies spatial location and molecular signature of perivascular niche as prognostic covariates

We sought to further investigate the spatial heterogeneity of vasculature organizations. Inspired by the concept of geographic diversification of tissue subregions^33^, we proposed **STAMPER (S**patial **T**opology **A**nd **M**olecular **P**attern of p**ER**ivascular niche) – a metric to evaluate the correlation between molecular similarity and spatial proximity of perivasculatures, defined as the local environment extending up to 20 μm from each vessel’s boundary (**see Materials and Methods**). STAMPER assigned each tissue core to one of the two patterns: **clustered**, in which perivascular niches that share similar molecular profiles that tended to be spatially adjacent; or **random**, in which perivascular niches that share similar molecular profiles that were spatially independent (**Fig. 3D, E** and **Supplementary Figure S5D**). If no vessels were present, the tissue core was labeled as **depleted**. The random pattern was significantly associated with poor RFS compared to the depleted pattern (log-rank p = 0.039, **Fig. 3F**), with similar trends for OS. This finding is in line with our previous results that higher VD is a worse prognostic parameter.

### Survival group associated with distinct cellular infiltration profiles

To gain further insight into the impact of the tumor vasculature on local immune response, we profiled the dynamics of cell phenotype fractions as functions of distance to nearest vessel boundaries. We observed distinct cell infiltration profiles between the survival groups. Long-term survival was associated with significantly increased infiltration of total immune cells and T lymphocytes (**Fig. 3G**). Specifically, the significance of elevated infiltration was mostly associated with immune cells in long-term survivors, except that antigen-presenting cells (APCs) were more prevalent in short-term survivors. This aligns with the observation that tumor-infiltrating APCs may be associated with poor breast cancer prognosis^34^. Stromal cells were also more actively in infiltrating in long-term survivors, while tumor cells exhibited similar infiltration patterns across survival groups. This observation suggests that stromal cells might develop alternative mechanisms to favor the migration of immune cells than tumor cells for an enhanced antitumor immunity (**Fig. 3H**). To gain further insights into the interactions between different cell populations, we tested the correlation between each pair of infiltration profiles using Pearson correlation method and summarized in a heatmap. Consistent with analyses in breast tumors^35^, tumor cells had a universal tendency towards anti-correlation with immune cells in long-term survivors. Conversely, tendencies towards positive correlations were identified with Treg and Tex cells, B cells, CD57^+^ cells, and macrophages in short-term survivors. Of note, a meta-cluster of macrophages and granulocytes was positively correlated with effector lymphocytes in short-term survivors (**Fig. 3I, box a-c**), suggestive of a possible myeloid-related immune suppression^36–39^.

### Spatial analyses identified characteristics of cellular neighborhoods that are correlated with survival

Our results suggested that immune response in the TNBC TME was coordinated by spatially-resolved cellular interactions. To assess this, we followed a canonical approach to construct cellular neighborhoods by first identifying the top 20 nearest neighbors of each cell^40^ (**Fig. 4A**). We then clustered all cells into 10 distinct cellular neighborhoods (CNs), each characterized by the unique composition of cell phenotypes, and captured by local tissue architectures. On the basis of biological functions of the predominant cell phenotypes, CNs were annotated as follows: tumor compartment (CN1), tertiary lymphoid structure-like (CN2), immune-fibroblasts interface (CN3), granulocyte-enriched (CN4), pan-immune hotspot (CN5), Ki67^+^ immune hotspot (CN6), CD57^+^-enriched (CN7), myofibroblast-enriched (CN8), Treg-enriched tumor (CN9) and perivascular immune niche (CN10) (**Fig. 4B**). To assess the survival associations of CNs, we compared the CN compositions across survival groups and revealed significantly more CN5 cells in long-term survivors (Wilcoxon rank-sum *p* = 0.0289). Using the median percentage of cells for each as the cutoff threshold, we confirmed this association between CN5 and prolonged survival using Kaplan-Meier estimates. Of note, CN4 and CN10 were also independently associated with improved survival. These findings add to our previous results that while the presence of general vasculature niches might be linked to poor survival, perivascular immune niches were associated with improved survival, suggesting an anti-tumor role (**Fig. 4D**, **Supplementary Figure S6**).

**Figure 4.**
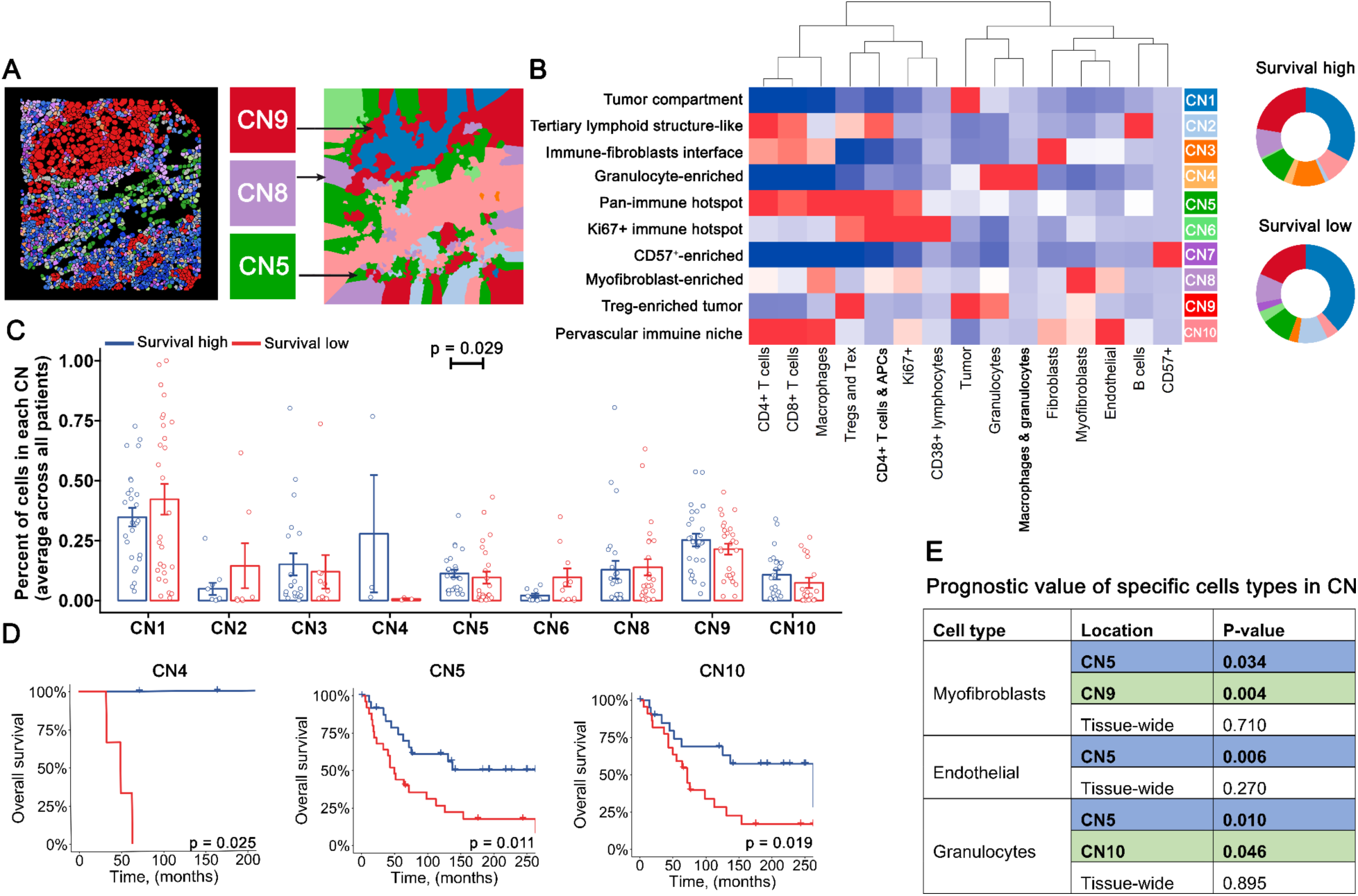
Spatial neighborhood analyses. **(A)** Analytical workflow for identification of cellular communities based on top 20 nearest neighbors of each cell. **(B)** Cells were clustered into 10 cellular neighborhoods (CN) that recapitulate unique local tissue architectures. **(C)** Prevalence of CNs across survival groups as percentage of cells. **(D)** OS plots for CN prevalence where patients were classified into high or low group based on the median of CN frequencies. P-values were computed using log-rank test. **(E)** Prognostic value in relation to overall survival of specific cell types in CNs. P-values were computed using Cox proportional hazard ratio model.

We next applied the Cox proportional hazard ratio model to determine the survival significance of individual cell phenotypes within each CN. The frequencies of myofibroblasts and granulocytes in CN5 and CN9 and endothelial cells in CN5 were significant prognostic factors, suggesting that these cells could be modulating antitumor immunity in a region-specific manner (**Fig. 4E**).

We next sought to quantify the interactions between CNs and their relationship to clinical outcomes. We developed a network-based algorithm to transform spatial cell maps into connected graphs, in which each cell represents a node and edges represent spatial proximity. For each CN, the algorithm extracted cells that connected to cells from other CN types (**Fig. 5A**). We next quantified the absolute number of border cells for each pair of CNs and partitioned them into long-term and short-term survival groups. Numbers in long-term group were subtracted from short-term group to reflect the differential CN interactions across survival groups (**Fig. 5B**). Of note, we showed that Treg-enriched CN (CN9) interacted more with tumor CN (CN1) but less with myofibroblast CN (CN8) in long-term survivors. Focusing onto borders between CN1-CN9 and CN8-CN9, we observed an enrichment of CD8^+^ T cells in long-term survivors, which suggests the CD8^+^ T cell activity at specific CN borders could be important in the antitumoral immune response (**Fig. 5C**). Assuming the major cell type of CNs as the phenotypic cell type, we adapted a canonical proxy to model spatial cellular interactions as a simplified surrogate for CN interactions, denoted as **S**patial regulatory and exhausted **T** cell **A**ffinity **IN**dex (**STAIN**) (**Fig. 5D**). We computed the distances of each regulatory or exhausted T cell (defined as Treg and Tex in the original source, represent CN9) to its nearest tumor cell (*d*_1_, represent CN1) and myofibroblast (*d*_2_, represent CN8). **STAIN** is thus defined as *d*_1_/(*d*_1_ + *d*_2_), which reflects the affinity of Treg and Tex cells towards tumor cells versus myofibroblasts (**Fig. 5E**). The Treg and Tex cells were significantly more adjacent to tumor cells in long-term survivors, which recapitulated the high interaction score between CN1-CN9 (**Fig. 5F**). Importantly, we observed the same trend in the validation cohort. The patients with pathological complete response (pCR, responders) after neoadjuvant combination chemotherapy with or without atezolizumab had significantly lower STAIN (i.e., lower relative spatial proximity of Treg and Tex cells to tumor cells) at baseline compared to patients with residual disease (RD, non-responder), suggesting that STAIN could also serve as a predictive biomarker (**Fig. 5G**). In addition, exclusion tests were performed for both cohorts such that patients were excluded one at a time and the same significance was still observed among the remaining patients, suggesting the robustness of STAIN (**Supplementary Fig. S7**).

**Figure 5.**
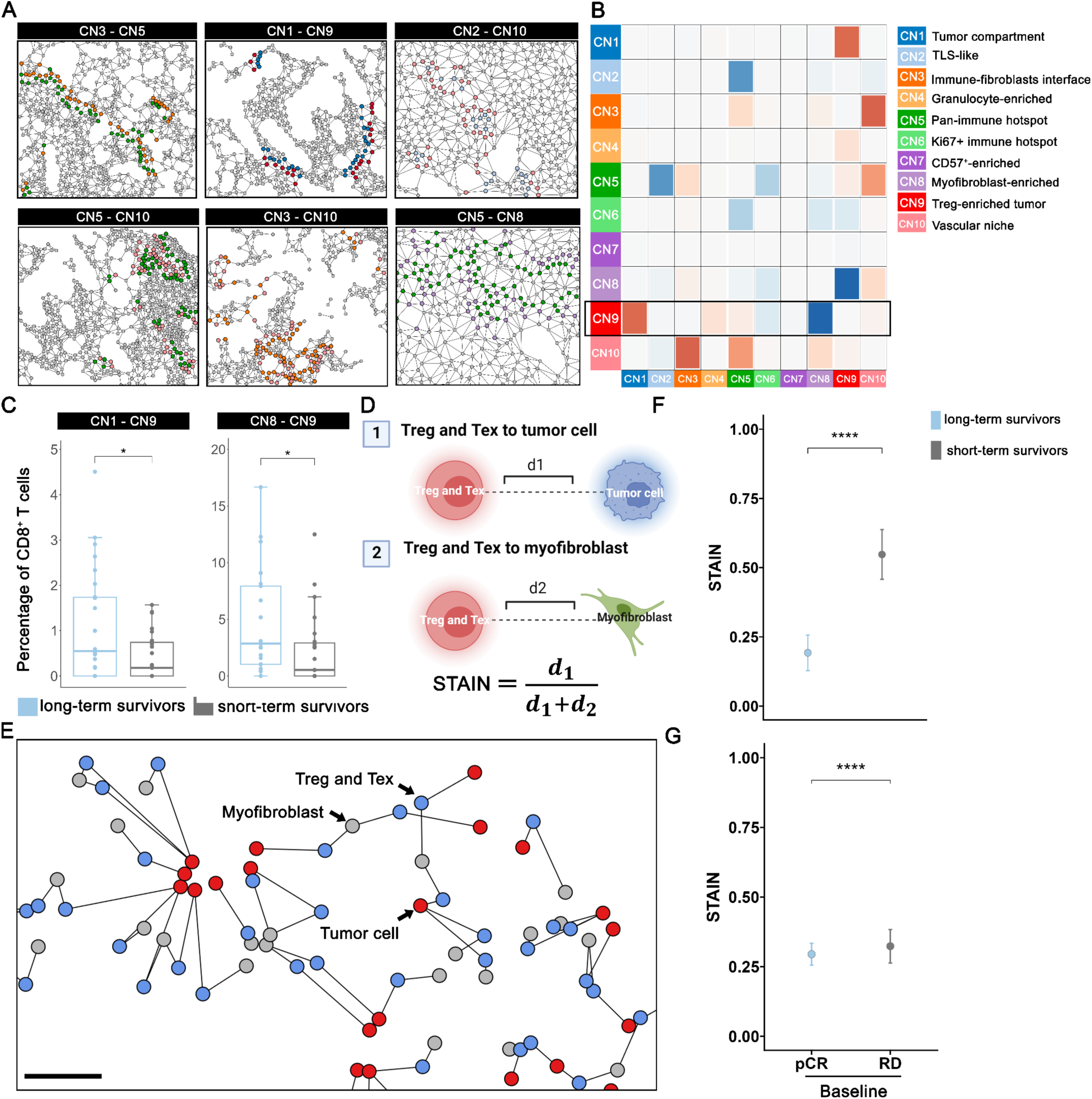
Interactions between cellular neighborhoods are associated with clinical outcome and predictive of therapeutic response to cancer treatment. **(A)** Representative subregions of cellular neighborhood (CN) interactions. Connections between cells of different types of CN were considered as interactions. **(B)** CN interaction heatmap that reflected the difference in interaction intensities between survival groups. Brown: high intensity; blue: low intensity. **(C)** Prevalence of CD8^+^ T cells at CN1-CN9 and CN8-CN9 as percentage of cells. **(D)** Mechanistic model of spatial tumor cell affinity index (STAIN), derived from the relative distances from tumor cell to its nearest Treg and Tex cells and myofibroblast. **(E)** Representative topological graph of tumor cells, Treg and Tex cells and myofibroblasts within a given tumor. Scale bar, 50 μm. STAIN was computed on a per-cell basis and compared between survival groups. **(F)** For validation, STAIN was computed on a per-cell basis for baseline and post-treatment samples and compared between patients with pathologic complete response (pCR, responders) and patients with residual disease (RD, non-responders) after neoadjuvant chemotherapy with or without atezolizumab. Statistical comparisons were performed using Wilcoxon rank-sum test. ****, *p* < 0.0001; **, *p* < 0.01; *, *p* < 0.05.

### Predicting therapeutic response using deep learning

Given our results that engineered spatial cellular and tissue textural features are associated with patients’ survival, we wondered whether we could leverage a deep learning approach to minimally processed spatial data to predict clinical outcomes. To prepare the data, we generated spatial proximity graphs for both cells and vessels. These graphs were considered as coarse representations for cellular and tissue textures. 100 responders and 100 non-responders were randomly selected for analysis. Next, we utilized the FEATHER algorithm^41^ to generate one-dimensional embeddings for each graph, which were then concatenated and aggregated to patient level as the final feature matrix (see **Methods**). For proof of principle, we took advantage of an artificial neural network with canonical multi-layer perceptron architecture. Using the five-fold cross-validation method, we split the data into five partiotions, with four of the partitions (80% of the patients) as the training data and one of the partitions (20% of the patients) as the testing data to evaluate the prediction accuracy. This process was repeated five times to exhaust all possible combinations (**Fig. 6A**). We first tested the variations of feature matrices across response groups using principal component analysis on a cell graph-derived feature (CGD) matrix, a vessel graph-derived feature (VGD) matrix, and the combination of both. Results showed that responders and non-responders were well-blended regardless of feature categories, suggesting that it is challenging to effectively separate responders from non-responders with linear patterns (**Fig. 6B**). However, with deep learning, we showed that both CGD and VGD features could independently predict response to treatment, with average AUCs = 0.66 and 0.60 respectively (**Supplementary Table S2**). The combination of two feature sets with the same model further achieved an unprecedented average AUC = 0.71 (**Fig. 6C**). Comparison between AUCs from cross-validations indicates that CGD features have similar predictive power compared to VGD features (student t-test *p* = 0.20); however, their combinations significantly improved the prediction accuracy compared to each individual set (student t-test *p* = 0.04 and 0.02, respectively; **Fig. 6D**). Of note, our best model was able to enrich for non-responders, in which we observed mean specificity (true-negative rate) of 89% compared to the non-response rate of 50.9% in the NeoTRIP cohort. This suggests that deep learning with spatial data was especially capable of selecting patients that are not likely to respond to chemotherapy with or without neoadjuvant atezolizumab in the context of TNBC, thus avoiding any unnecessary immune toxicity.

**Figure 6.**
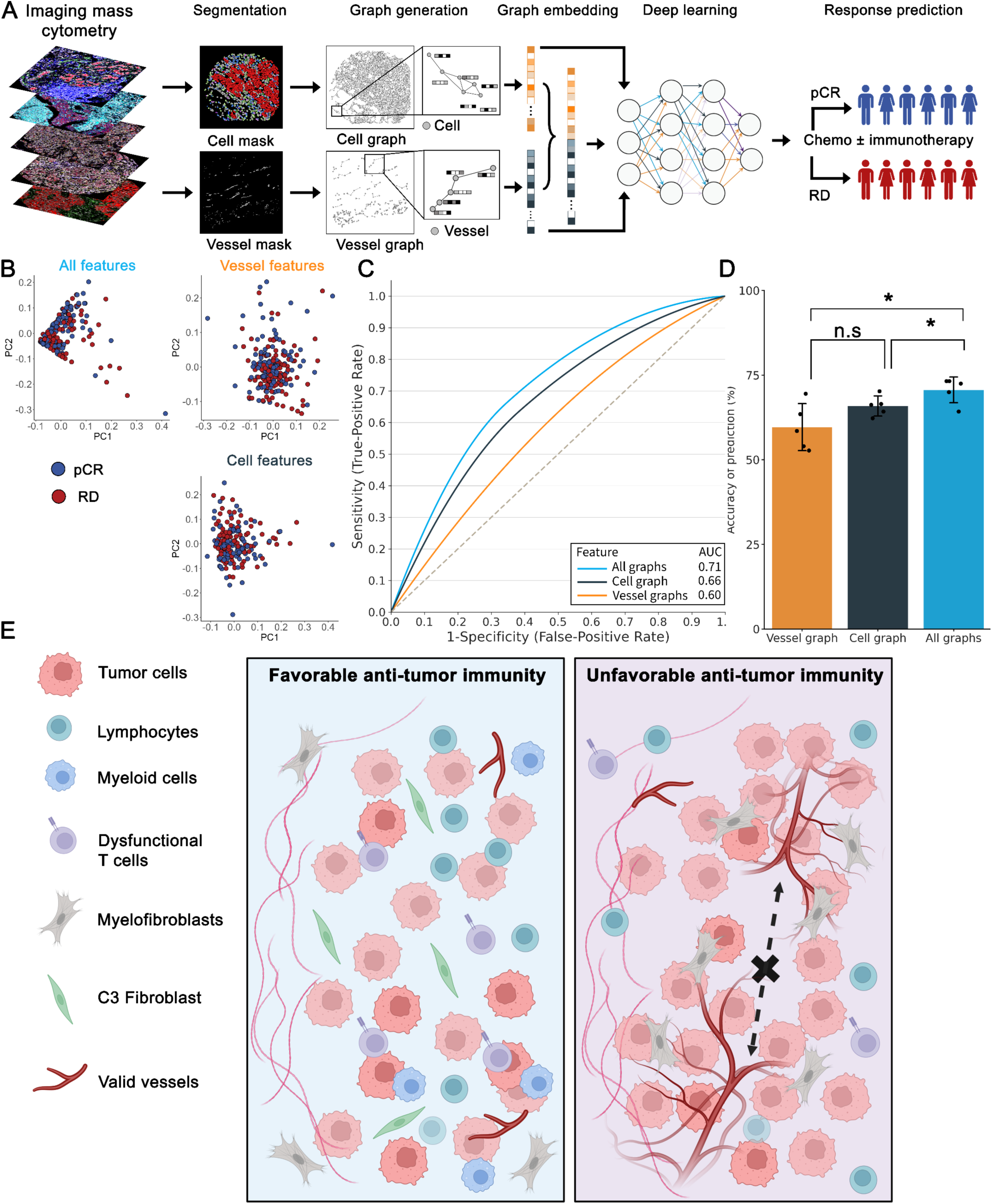
Deep learning of spatial data predicts therapeutic response in TNBC. **(A)** Analytical workflow for therapeutic response prediction using artificial neural network that was trained and tested on spatial cell and tissue data derived from imaging mass cytometry images. **(B)** Principal component analysis (PCA) on cell graph derived (CGD), vessel graph derived (VGD) features and their combination. **(C)** Smoothed (keep AUC fixed) receiver operating characteristics curve reflecting the mean prediction accuracy of the model with CGD, VGD and their combination as input using five-fold cross-validation. Mean AUCs were computed. **(D)** Comparisons between prediction accuracies obtained from three experiments with VDG feature alone, CDG features alone, and their combinations. Model with CDG features achieved similar accuracies compared to VDG. Their combinations yielded the highest mean AUC and was significantly higher compared to using individual set. **(E)** Conceptual framework for describing differential TME spatial parameters in TNBC with favorable or unfavorable anti-tumor immunity. Statistical comparisons, either paired or unpaired, were performed using student’s t-test. *, *p* < 0.05; n.s., non-significant, *p* > 0.05.

Taken together, our results have demonstrated that cellular and tissue dynamics in the TME are important predictors of treatment response in TNBC, which corroborates our hypothesis that spatial data are associated with clinical outcomes. We propose two mechanistic models of how the compositional and spatial behavior of the TME in clinical subgroups could be different in cell infiltration and topology, fibroblast subclustering, cellular neighborhood interactions and perivasculature profiles. In a favorable setting, the TME is marked by lymphocyte infiltration, low vascular density, and enrichment of a specific fibroblast cluster characterized by SMA expression. Conversely, in an unfavorable setting, the TME is infiltrated by myeloid cells and has higher vascular density, where vessels in proximity tend to be surrounded by perivasculature niches with similar molecular profiles (**Fig. 6E and Discussion**).

## DISCUSSION

This study advances understanding of immuno-oncology and clinical insights for patients with triple-negative breast cancer by using a multi-scale computational framework. Our approach captures the differential features of the tumor microenvironment as prognostic and predictive biomarkers for effective patient stratification. The ability to make such distinctions is helpful to inform the personalized care of patients predicted to respond to chemo/immuno-therapy or to identify optimal therapy for patients predicted to recur. The present study focuses on characterizing the tumor microenvironment organization with IMC of breast tumors. This approach, together with other spatial multiplex proteomics technologies options including OPAL multiplex immunohistochemistry (mIHC)^39,42^, co-detection by indexing (CODEX)^43,44^, multiplexed ion beam imaging (MIBI)^45^ and cyclic immunofluorescence (CyCIF)^46,47^, introduced a new paradigm for the initiating biomarker discovery at an unprecedented resolution. However, computational methods have unfortunately lagged behind the technological advancements^48^. At present, few engineering methods are capable of handling high-dimensional imaging data. Deep learning methods are competitive but often sacrifice interpretability to users with limited computational backgrounds^49^. By integrating quantitative tissue profiling, multi-scale spatial analyses and deep learning with interpretable sources, this study aids in the advancement of the computational framework as an efficient, informative tool in biomarker discovery.

In this study, we take advantage of two public single-cell datasets from breast tumors for method development and validation. A primary cohort of 58 patients with TNBC (discovery cohort) from the METABRIC study was selected. The corresponding single-cell database of 131,001 objects feaured various subtypes of lymphoid cells, myeloid cells, tumor cells and stromal cells and their functional states. The methods described here represent a hierarchical analysis of the TNBC TME landscape spanning single-cell level properties and tissue-level patterns. In addition to the TNBC specimens evaluated herein, the backbone of this computational suite has been used for biomarker discovery in the context of bladder cancer with sequential IHC imaging^50^, hepatocellular carcinoma with IMC and pancreatic cancer with mIHC^38^, therefore providing a general digital pathology infrastructure for immuno-oncology.

The overarching goal of the methods was to discern differential biological behaviors of tumors in patients with poor and superior clinical outcomes for the identification of potential biomarkers. To start, we quantified the overall TME architecture by computing the distribution of each cell population and observed great heterogeneity in tumor-immune compositions. Despite great variance in densities, immune infiltrations were coordinated as we observed significant positive correlations between immune cell densities with B cells but negative correlations with macrophages. Next, we demonstrated that cell distributions were linked to various clinical outcomes, including TMB level, age of diagnosis, tumor grade and primary tumor laterality.

Importantly, we further observed a prevalence of B cells and CD4^+^ T cells in long-term survivors in line with what is known about anti-tumor immunity in TNBC. In addition, we identified that fibroblasts were prevalent in long-term survivors. Subclustering on fibroblasts based on expression of four canonical fibroblast markers - SMA, Caveolin-1, PDGFRB and FSP1 showed that a specific fibroblast subpopulation with a profile of SMA^Hi^ Caveolin-1^Low^ PDGFRB^Low^ FSP1^Neg-Low^ was elevated in long-term survivors. Despite the prognostic role of fibroblasts as presented in this study, conflicting information on the association of fibroblasts and clinical outcomes has been suggested for breast cancer and pancreatic cancer^51–55^. Thus, further investigation evaluating the precise molecular phenotyping of fibroblasts to fully elucidate their roles in immuno-modulation are warranted.

Using vasculature masks, we first demonstrated that a high valid vasculature density (area greater than 500 μm^2^) was associated with poor RFS. To further understand the spatial and functional implications of the tumor-associated vasculature, we proposed STAMPER, namely spatial topography and molecular pattern of perivascular niche. STAMPER determined that tumors with randomly distributed vessels were associated with poor RFS relative to those without vessels. Associated with the vascular heterogeneity, we detected great variations in cell infiltration profiles among survival groups. Correlation analyses showed that synergistic or antagonistic effects between infiltration patterns of different cell types may explain the differential clinical outcomes. Such correlations motivated us to further interrogate the cellular interactions. In this study, we adopted a canonical approach to define cellular neighborhoods (CNs). We showed that enrichment of pan-immune CN and perivascular immune niche CN were coupled with increased interactions between Treg and Tex cell-enriched tumor CN and the tumor compartment in long-term survivors. We further proposed a mathematical metric to model such interaction, namely STAIN (spatial regulatory and exhausted T cell affinity index), which recapitulates the positive link between tumor cell proximity to Treg and Tex cells and improved survival. The predictive value of STAIN was corroborated in the validation set, a cohort of TNBC patients recruited to the NeoTRIP clinical trial and treated with chemotherapy alone or combined with immunotherapy, as responders were associated with lower STAIN. Nevertheless, such results shall be interpreted with caution, considering the known immunosuppressive interplay between tumor cells and Treg and Tex cells such as regulatory T cells.

Finally, we trained deep learning models using engineered spatial data from NeoTRIP cohort to predict response to chemotherapy versus chemo-immunotherapy. Specifically, we constructed two sets of graphs that represent cellular and tissue textures. Embeddings from these graphs were then computed as features for prediction. We showed that spatial cellular and tissue textures can independently predict response to therapy. The combination of two feature sets further yielded notable accuracy a mean of AUC of 0.71 with an enriched non-response rate of 89% in predicted non-responders relative to baseline non-response rate of 51%. Such findings represent an important advance in reducing the unnecessary toxicity of cancer treatment. It is also worth noting that the deep learning model was trained on biologically meaningful features, which preserved both the interpretability and efficiency compared to existing prediction tools.

In conclusion, we developed a multi-scale framework using two public TNBC datasets and identified key spatial and compositional biomarkers that are predictive of clinical outcomes. The framework represents a conceptual infrastructure and important computational solution for analyzing high-dimensional imaging data in search of biological cues underlying human diseases. The framework, when expanded to larger, well-annotated patient cohorts, may provide the insights needed to develop informed therapeutic clinical trial designs, resulting in transformative and personalized clinical interventions^56^. The data obtained in this study will also be very valuable for spatial computational quantitative systems pharmacology modeling to advance precision oncology^57^.

## MATERIALS AND METHODS

### Dataset Acquisition and Processing

Two spatially resolved single-cell datasets derived from triple-negative breast cancer (TNBC) patients were obtained from public data repositories (see **Data availability**). The primary dataset was a subset of the METABRIC study (*n* = 718) that contains only TNBC subtype (*n* = 58) and was used for method development and biomarker discovery. As described in the original METABRIC study^13^, ER, PR, and HER2 status were classified based on their empirical expression distributions using a 2-component Gaussian mixture model as implemented in MCLUST^58,59^ in an approach similar to that described by Lehmann and colleagues^60^. The secondary dataset contains previously evaluated patients in a randomized phase 3 clinical trial (NeoTRIP)^17^ with chemotherapy or without immune checkpoint blockade therapy (anti-PD-L1, atezolizumab), with tumor sampled at three time points (baseline, *n* = 243; early on-treatment, *n* = 207; post-treatment, *n* = 210). A subset containing baseline and post-treatment samples were subjected for biomarker validations. Clinical characteristics are available in respective sources. Both datasets are presented in processed format, where single-cell measurements including spatial coordinates and phenotype annotations are available. For consistency, coordinates are transformed from pixels to microns using the resolution provided by the respective imaging modality. In both datasets, cells with the original label “Epithelial” were merged into a master label “Tumor” (as also defined in the original study) as we did not further distinguish epithelial subpopulations in this study. Similarly, “myofibroblasts PDPN+” cells are merged with “myofibroblast” cells and “fibroblasts FSP1+” were merged with “fibroblasts” cells in the discovery dataset.

### Spatial topology and molecular pattern of perivascular niche (STAMPER)

Vessels with area greater than 500 μm^2^ (valid vessels) were subjected to assessment using STAMPER. For each valid vessels, cells within 20 μm were identified (perivascular niche) and their protein expressions were averaged to formulate a molecular profile at per-vessel basis. For each pair of valid vessels within the same tissue core, Euclidean distance between molecular profile vectors was computed and denoted as “molecular distance”, and Euclidean distance between the center of cells were computed and denoted as “geographical distance”. Mantel test was then performed to assess the correlation between molecular and geographical distances. Cores with 2 or more valid vessels were labeled as either “clustered” (Mantel *p* < 0.05) or “random” (Mantel *p* >= 0.05). Cores without valid vessels were labeled as “depleted”.

### Cellular neighborhood identification

A “window” capturing the top 20 nearest neighbors (cells) was identified for each cell using C++ accelerated kd-tree algorithm, which was implemented using “nn2” function from R package “RANN”. Each “window” was then transformed into a vector containing the frequency of each cell type. Cells with their defining “windows” were then clustered into 10 groups using unsupervised hierarchical clustering with MiniBatchKMeans algorithm with batch size = 100 from R package “ClusterR”. Based on the majority cell types, each group was then annotated with a biological phenotype to formulate the final cellular neighborhood.

### Cellular neighborhood interactions

Cells that established heterotypic connections were considered as interacting cells between cellular neighborhoods (CNs). The cumulative number of such connections was regarded as the communication intensity between each CN. A spatial network-based algorithm was developed to quantify CN communications. For each tissue core, Delaunay triangulation was generated and long edges (> 30 μm) were assumed to have low impact on connected cells and were removed. The remaining graph was then analyzed to locate connections that linked heterotypic cells for each pair of CNs and the number of such connections were denoted as interactions.

### Spatial regulatory and exhausted T cell affinity index (STAIN)

For each tumor cell, we computed the distance to its nearest regulatory and exhausted T cell (cell type Treg and Tex as described in the original source^13^), denoted as *d*_1_; and the distance to its nearest myofibroblast, denoted as *d*_2_. STAIN is therefore defined as:

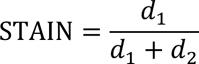

STAIN is designed to recapitulate the proximity between tumor compartment (CN1) and Treg-enriched tumor (CN9) and avoidance between CN1 and myofibroblast CN (CN8). As there was no “Treg and Tex” cell type in the validation cohort (NeoTRIP cohort)^16^, we merged “Tregs” and “CD8^+^PD1^+^ Tex” cell types as an approximation.

### Deep learning

All deep-learning analysis steps were performed in Python (version 3.10.12). We used the TensorFlow (version 2.13.0) framework with Keras serving as an interface. The neural network model was built using a sequential architecture. Hyper-parameters, including optimizer, loss function and learning rate were optimized using the package optuna (version 3.4.0). To prepare data for training and testing, we engineered two sets of graphs based on cell and vessel topography. For each sample (tissue core), we first generated Delaunay triangulation for all cells and removed links greater than 30 µm to formulate the cell graph. Each node (cell) in the graph was coded with a feature vector comprising of cell type, centroid coordinates and neighbor compositions (**Supplementary Table 1**). Next, we connected all vessels with their nearest neighboring vessel to formulate the vessel graph. Each node (vessel) in the graph was coded with a feature vector comprising of molecular profiles, that is, the averaged proteomic expressions of all cells that within 20 µm to the vessel boundary. We then used FEATHER algorithm to extract graph embeddings (length = 250) from each graph to formulate two sets of features: cellular graph derived (CDG) and vessel graph derived (VDG) features. For each set, features were then aggregated from sample (core) level to patient level by taking the mean, maximum and minimum to construct final feature matrix (length = 750).

For training and testing, we randomly selected 100 patients each from both responders and non-responders group using sample function with random_state parameter set to 1 for reproducibility. Five-fold cross-validation method was used to evaluate the prediction accuracy. Here, we have three modes of data for our experiments: (1) CDG features only; (2) VDG features only and (3) combination of two sets. Mean area under the receiver operating characteristics (AUC) curve after cross-validations was computed as the measurement of prediction accuracy. Smoothed ROC curves were plot for comparison. Various classes of Scikit-learn (version 1.2.2) machine-learning libraries have been utilized for the training and testing tasks. Graph generations and node feature encoding were performed using R. Graph embedding extractions and training steps were performed on Google Colab GPU/TPU servers. See the ‘Code availability’ section for additional details.

## Statistical analysis

Wilcoxon rank-sum test was performed for comparisons between paired and unpaired data using “Wilcox.test” function from R package “stats”. Student’s t test is used to compare the mean AUCs between experiments for deep learning model evaluations. Bonferroni correction was used for multiple tests *p* values using “p.adjust” from R package “stats” and adjusted *p* < 0.05 was considered significant. Pearson correlation coefficients were computed to assess correlation between two sets of data using “cor” function from R package “stats”. OS and RFS were estimated using Kaplan Meier method and statistical differences were assessed using log-rank tests using “survfit” function from R package “survival”. Cox proportional hazard ratio model was used to assess the prognostic value of cell types in specific CNs using “coxph” function from R package “survival”. Mantel test was performed using “mantel” function from R package “vegan”.

## Data availability

Single-cell dataset and clinical information supporting the biomarker discovery are available at the Zenodo data repository (https://zenodo.org/record/5850952). Single-cell dataset and clinical information supporting biomarker validations are available at (https://zenodo.org/record/7990870) for academic non-commercial research. For commercial access to the secondary dataset, parties will be directed to appropriate contact as declared in the original study.

## Code availability

For the primary cohort, codes for IMC image processing are hosted at https://github.com/BodenmillerGroup/ImcSegmentationPipeline, https://github.com/CellProfiler, and https://github.com/ilastik. Cell clustering were conducted using R packages FlowSOM (https://github.com/saeyslab/FlowSOM) and Phenograph (https://github.com/JinmiaoChenLab/Rphenograph). For the secondary cohort, codes for IMC image processing is hosted at https://github.com/BodenmillerGroup/ImcSegmentationPipeline and https://github.com/vanvalenlab/deepcell-tf. All additional analysis code that is original in this study can be accessed https://github.com/Shawnmhy/Spatial_predictors_TNBC.

