## Supplemental Files for "Spatial and Compositional Biomarkers in Tumor Microenvironment Predicts Clinical Outcomes in Triple-Negative Breast Cancer"

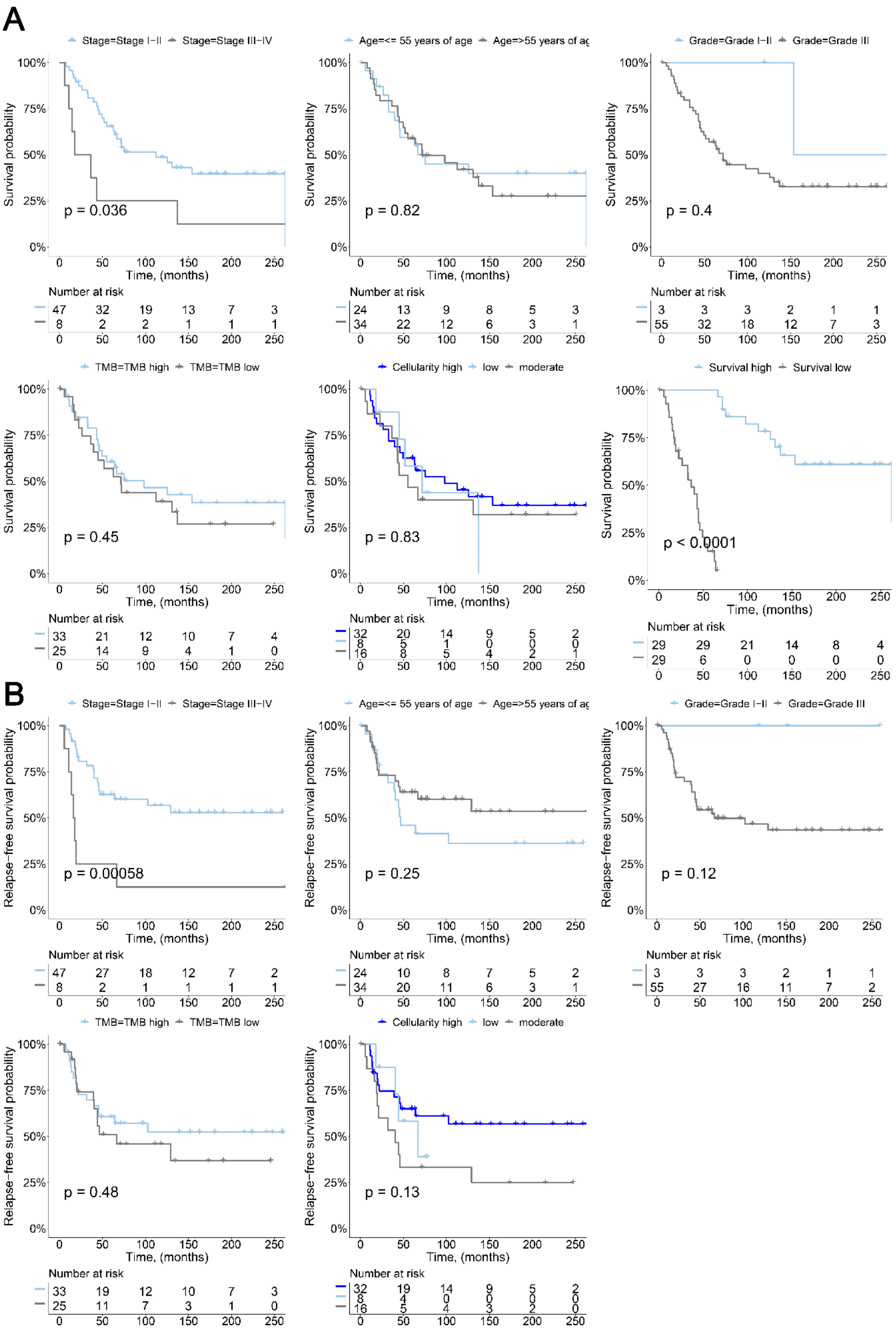
A
Stage=Stage I−II Stage=Stage III−IV
Survival probability
p = 0.036
Time, (months)
Number at risk
47 32 19 13 7 3
8 2 2 1 1 1
Age=<= 55 years of age Age=>55 years of ag
p = 0.82
24 13 9 8 5 3
34 22 12 6 3 1
Grade=Grade I−II Grade=Grade III
p = 0.4
3 3 3 2 1 1
55 32 18 12 7 3
TMB=TMB high TMB=TMB low
p = 0.45
33 21 12 10 7 4
25 14 9 4 1 0
Cellularity high low moderate
p = 0.83
32 20 14 9 5 2
8 5 1 0 0 0
16 8 5 4 2 1
Survival high Survival low
p < 0.0001
29 29 21 14 8 4
29 6 0 0 0 0
B
Stage=Stage I−II Stage=Stage III−IV
Relapse−free survival probability
p = 0.00058
47 27 18 12 7 2
8 2 1 1 1 1
Age=<= 55 years of age Age=>55 years of ag
p = 0.25
24 10 8 7 5 2
34 20 11 6 3 1
Grade=Grade I−II Grade=Grade III
p = 0.12
3 3 3 2 1 1
55 27 16 11 7 2
TMB=TMB high TMB=TMB low
p = 0.48
33 19 12 10 7 3
25 11 7 3 1 0
Cellularity high low moderate
p = 0.13
32 19 14 9 5 2
8 4 0 0 0 0
16 5 4 3 2 0

**Figure S1. Kaplan-Meier plots showed differential (A) overall survival and (B) relapse-free survival across clinical subgroups.**
Statistical analysis (log-rank test)

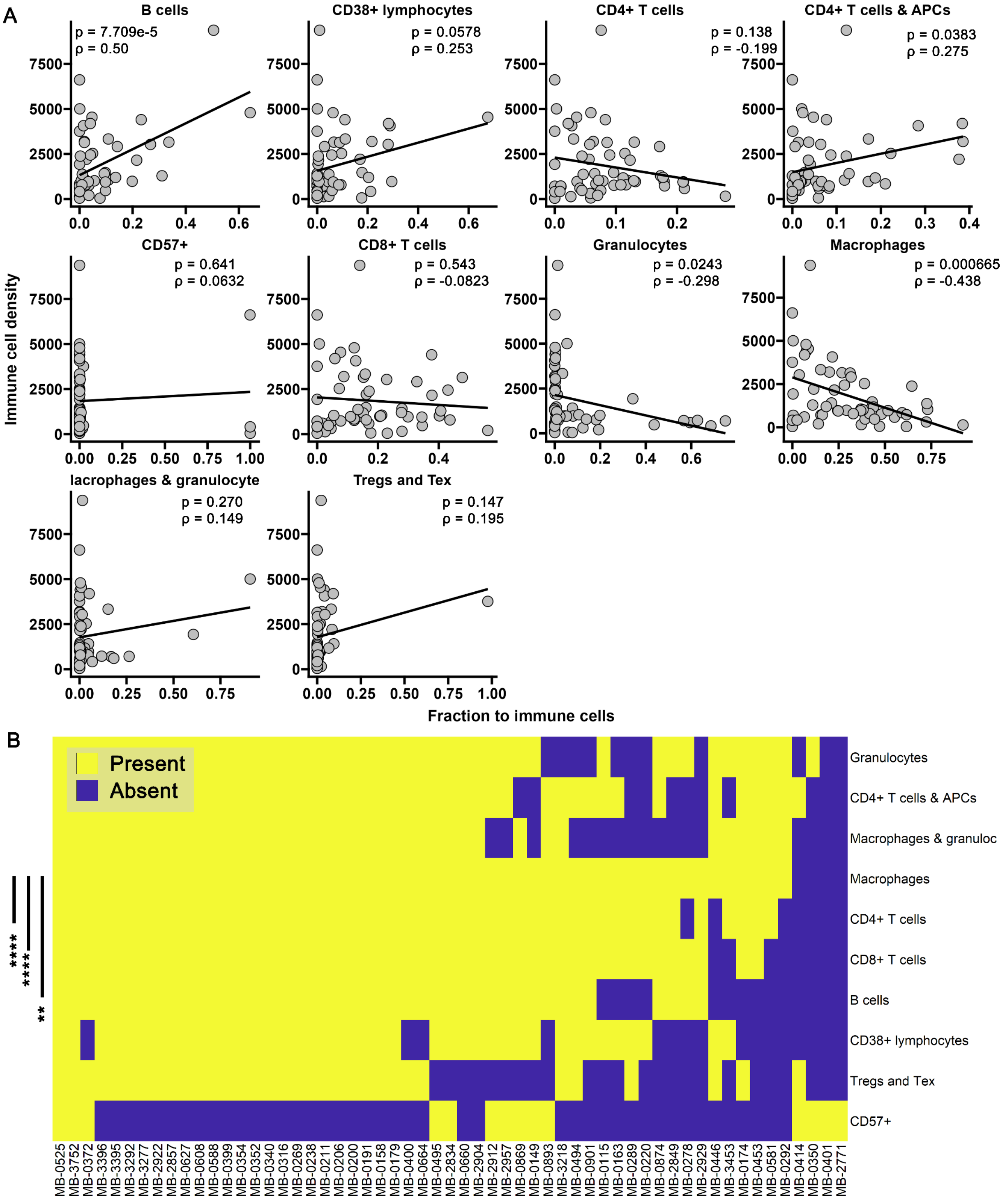


**Figure S2. Prevalence of immune subpopulations.**

(A) Correlation analyses between overall immune cell density and fraction to immune cells of each subpopulations.

(B) Presence versus absence analyses of immune subpopulations across patients.

Statistical analysis (A: Pearson correlation test)

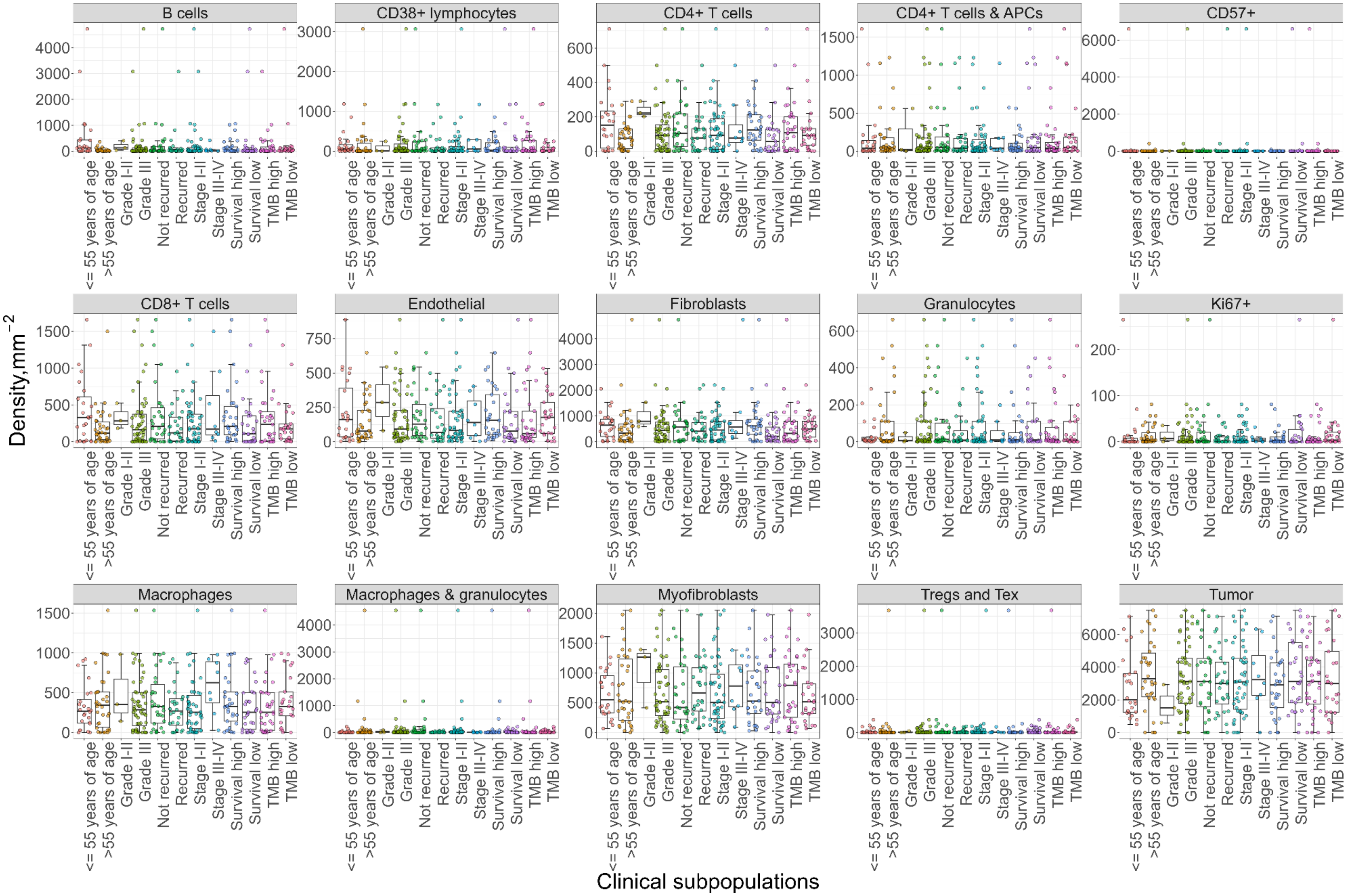


**Figure S3.** Prevalence of cell distributions across clinical subgroups as density.

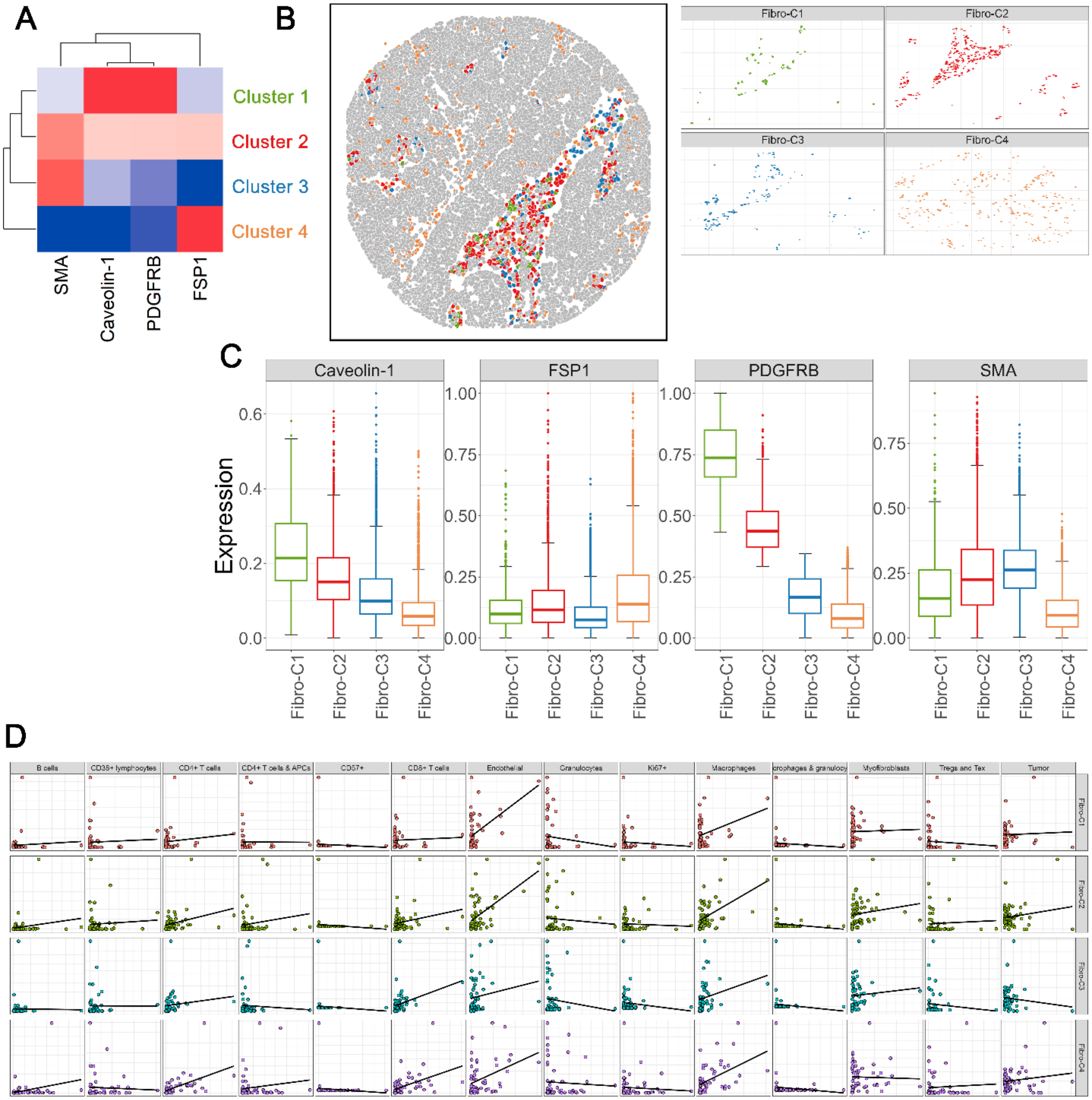


**Figure S4. Subclustering based on Caveolin-1, FSP1, PDGFRB and SMA expressions revealed compositional heterogeneity in TNBC patients.**

(A) Unsupervised hierarchical clustering on fibroblasts based on compositions of four markers: Caveolin-1, FSP1, PDGFRB and SMA.

(B) Representative spatial distributions of four fibroblast subclusters.

(C) Comparisons of four markers across four fibroblast subclusters.

(D) Correlation analyses between each fibroblast subcluster and other cell types.

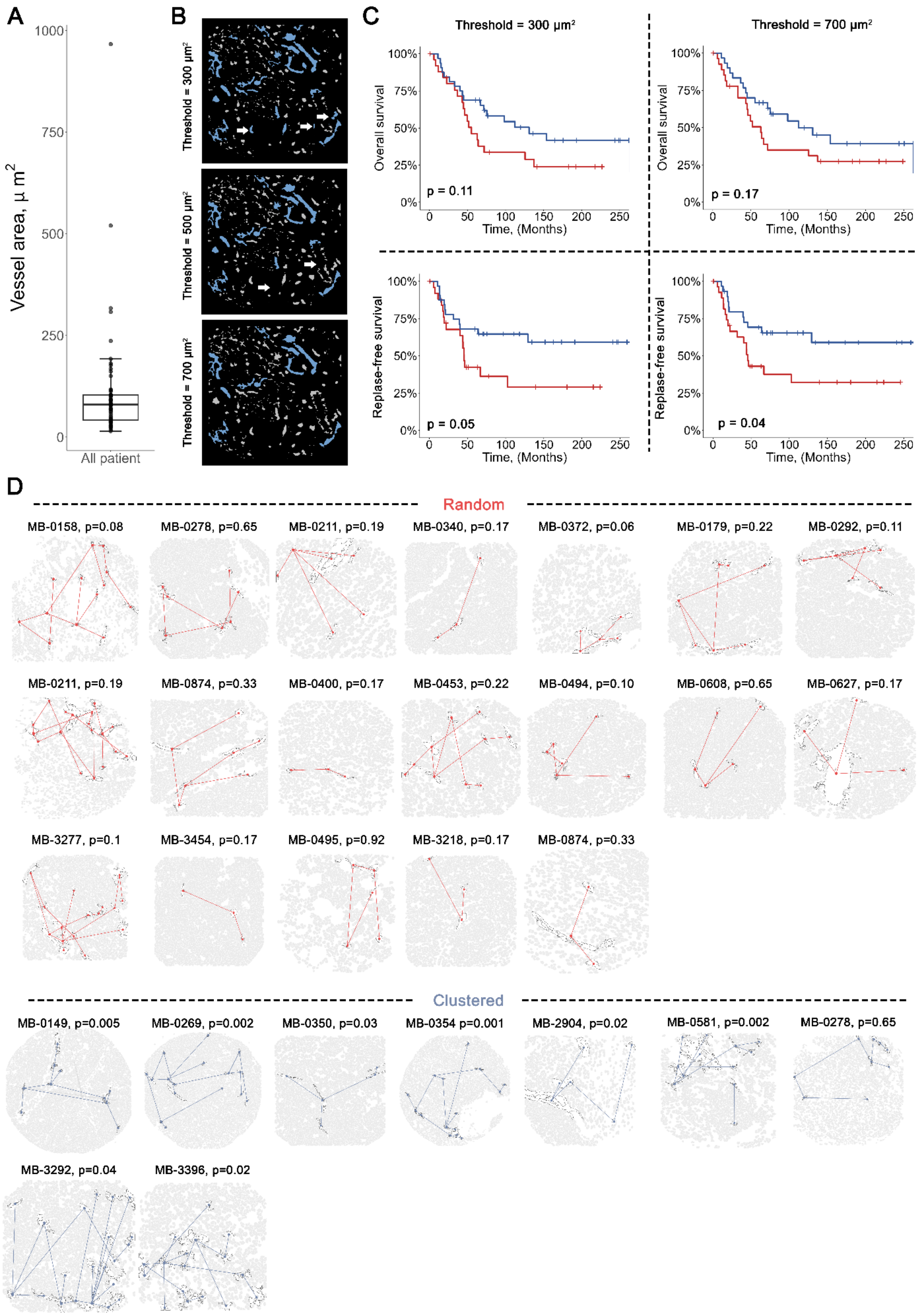

A
B
C
D
Vessel area, μ m2
All patient
Threshold = 300 μm²
Threshold = 500 μm²
Threshold = 700 μm²
Overall survival
Replase-free survival
Time, (Months)
p = 0.11
p = 0.17
p = 0.05
p = 0.04
Random
MB-0158, p=0.08
MB-0278, p=0.65
MB-0211, p=0.19
MB-0340, p=0.17
MB-0372, p=0.06
MB-0179, p=0.22
MB-0292, p=0.11
MB-0211, p=0.19
MB-0874, p=0.33
MB-0400, p=0.17
MB-0453, p=0.22
MB-0494, p=0.10
MB-0608, p=0.65
MB-0627, p=0.17
MB-3277, p=0.1
MB-3454, p=0.17
MB-0495, p=0.92
MB-3218, p=0.17
MB-0874, p=0.33
Clustered
MB-0149, p=0.005
MB-0269, p=0.002
MB-0350, p=0.03
MB-0354 p=0.001
MB-2904, p=0.02
MB-0581, p=0.002
MB-0278, p=0.65
MB-3292, p=0.04
MB-3396, p=0.02

**Figure S5. Prognostic value of valid vessel densities with various area thresholds**
(A) Prevalence of vessel areas averaged for each patient across the discovery cohort.
(B) Representative valid versus invalid vessels at different area thresholds. Threshold = 500 was selected for downstream analysis in the main text.
(C) Kaplan-Meier curves of overall and relapse-free survival for TNBC patients based on low and high valid vessel densities two alternative area thresholds confirmed that the prognostic value of valid vessels was robust.
(D) Visualizations of STAMPP algorithm results for all evaluable tissue samples.
Statistical analysis (B: Wilcoxon rank-sum test)

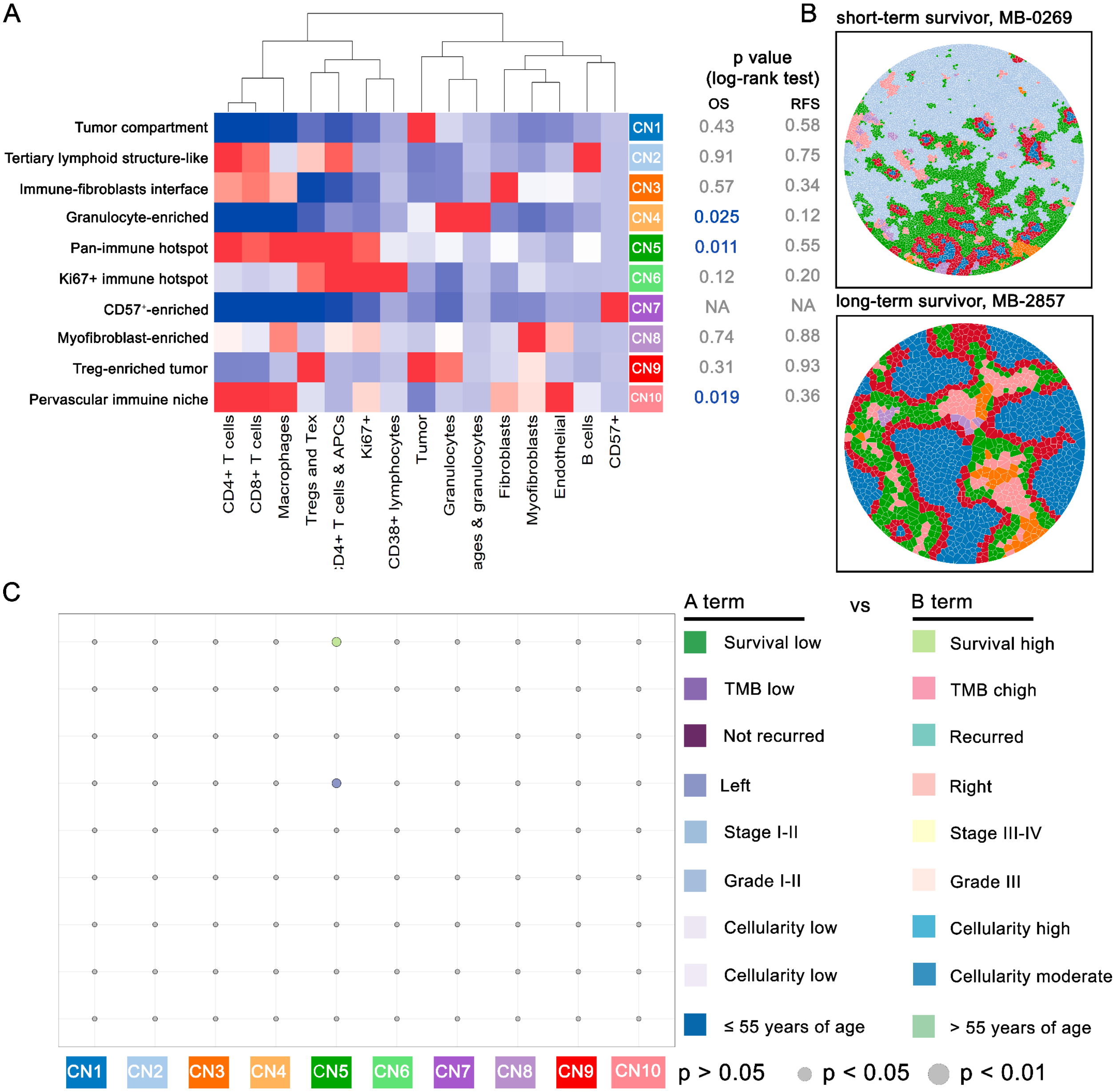


**Figure S6. Variability in 10 cellular neighborhoods (CNs) across clinical subgroups in TNBC**

(A) Heatmap of 10 CNs discovered in 58 TNBC patients.

(B) Representative tissues of 10 CNs using Voronoi diagrams.

(C) Bubble plot where circle color indicates which of the two terms listed on the right has higher levels of the CN type.

Statistical analysis (A: log-rank test; C: Wilcoxon rank-sum test)

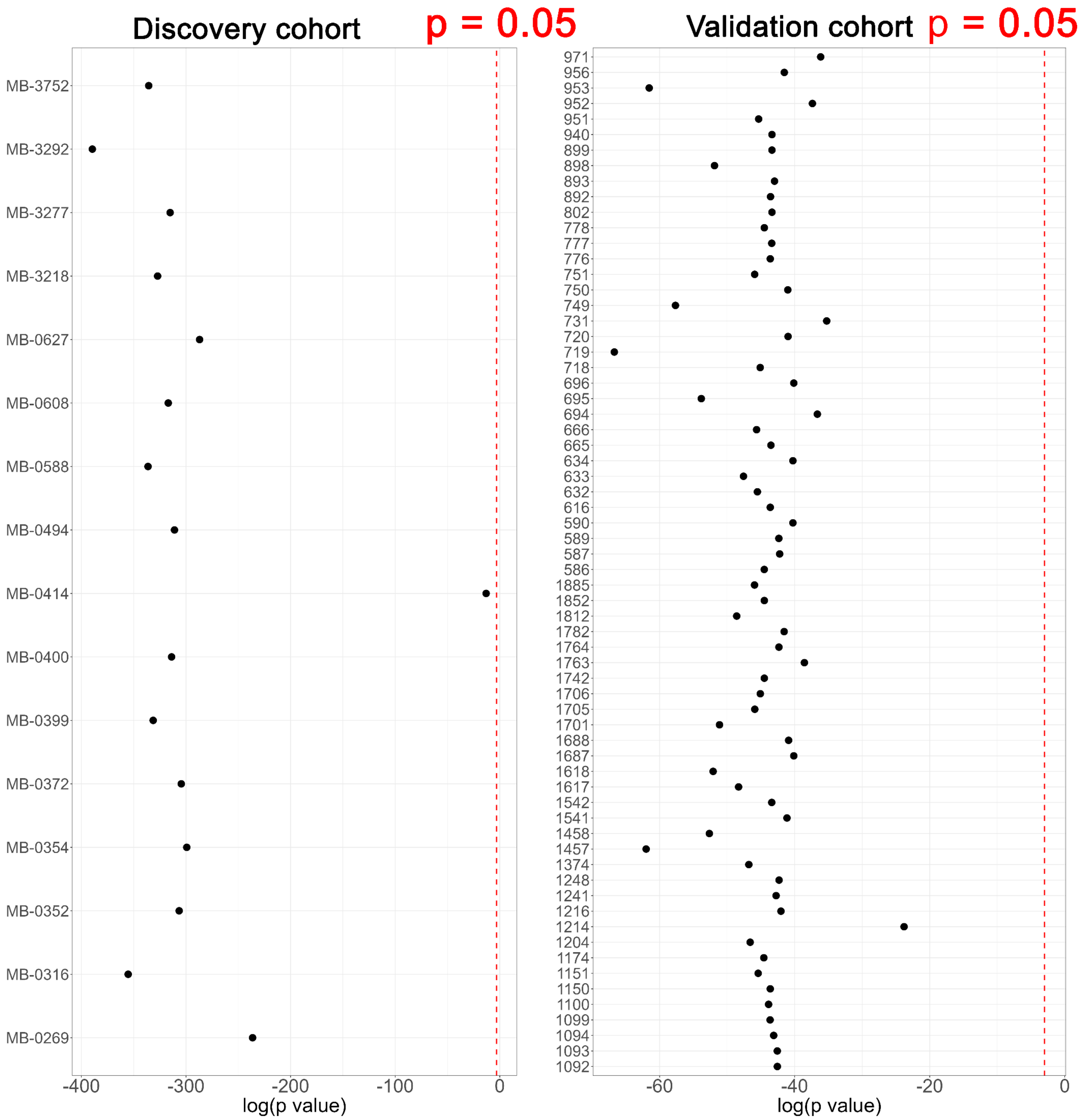


**Figure S7. Exclusion test to avoid systemic bias of STAIN introduced by a single patient on** (A) discovery cohort and (B) validation cohort.

Statistical analysis (Wilcoxon rank-sum test with holm method for multiple test comparisons)

| | |
|---|---|
| **Platform: Google Colab service** | |
| **Architecture: Multi-layer perceptron neural network** | |
| **Input layer** | |
| Dense layer | |
| No. neurons | 256 |
| Activation function | ReLU |
| Batch Normalization | |
| Dropout rate | 0.5 |
| **Hidden layer** | |
| **First – second hidden layers** | |
| Dense layer | |
| No. neurons | 256 |
| Activation function | ReLU |
| Batch Normalization | |
| Dropout rate | 0.5 |
| **Third to seventh layer** | |
| Dense layer | |
| No. neurons | 128 |
| Activation function | ReLU |
| Batch Normalization | |
| Dropout rate | 0.5 |
| **Output layer** | |
| Dense layer | |
| No. neuron | 1 |
| Activation function | Sigmoid |
| **Compilation** | |
| Optimization algorithm | Stochastic Gradient Descent (SGD) |
| Learning rate | 0.0928 |
| Loss function | Hinge |

Table S1. Configuration of deep learning algorithms

| Model performance | | |
| --- | --- | --- |
| **Settings for reproducibility** | | |
| Random seed for NumPy | 42 | |
| Random seed for TensorFlow | 42 | |
| Deterministic behavior for TensorFlow | Yes | |
| **Experiment 1** | | |
| **Configuration** | | |
| Feature set | Vessel graph derived (VDG) | |
| No. features | 750 | |
| No. responders selected | 100 | |
| No. nonresponders selected | 100 | |
| N-fold cross-validation | N = 5 | |
| Epochs | 800 | |
| **Evaluation** | **AUC** | **TNR** |
| Fold 1 | 0.635 | 0.90 |
| Fold 2 | 0.5275 | 0.80 |
| Fold 3 | 0.6950 | 0.95 |
| Fold 4 | 0.54 | 0.00 |
| Fold 5 | 0.585 | 0.90 |
| **Mean** | **0.60** | **0.71** |
| **Standard deviation** | **0.06** | **0.40** |
| **Experiment 2** | | |
| **Configuration** | | |
| Feature set | Cell graph derived (CDG) | |
| No. features | 750 | |
| No. responders selected | 100 | |
| No. nonresponders selected | 100 | |
| N-fold cross-validation | N = 5 | |
| Epochs | 800 | |
| **Evaluation** | **AUC** | **TNR** |
| Fold 1 | 0.6225 | 0.95 |
| Fold 2 | 0.6625 | 0.80 |
| Fold 3 | 0.6425 | 0.55 |
| Fold 4 | 0.7025 | 0.65 |
| Fold 5 | 0.605 | 0.85 |
| **Mean** | **0.66** | **0.76** |

| | | |
|---|---|---|
| **Standard deviation** | **0.03** | **0.03** |
| **Experiment 3** | | |
| **Configuration** | | |
| Feature set | CDG + VDG | |
| No. features | 750 + 750 | |
| No. responders selected | 100 | |
| No. nonresponders selected | 100 | |
| N-fold cross-validation | N = 5 | |
| Epochs | 800 | |
| **Evaluation** | **AUC** | **TNR** |
| Fold 1 | 0.7325 | 1.00 |
| Fold 2 | 0.725 | 0.90 |
| Fold 3 | 0.6425 | 0.85 |
| Fold 4 | 0.7325 | 0.80 |
| Fold 5 | 0.70 | 0.90 |
| **Mean** | **0.71** | **0.89** |
| **Standard deviation** | **0.03** | **0.07** |

Table S2. Configuration of training and testing and evaluations of model performance with different feature sets.
